# Oral sigma anti-bonding calcium carbonate promotes bone formation and delays myeloma progression in mice

**DOI:** 10.64898/2026.11.14.766415

**Authors:** Syed Hassan Mehdi, Dongjoon Lee, Alex Lee, Ha-Neui Kim, Junghwan Do, Maurizio Zangari, Fenghuang Zhan, Jong Y. Park, Paul K. Lee, Donghoon Yoon

## Abstract

Multiple myeloma is the most common cancer of the bone marrow and depends on the marrow niche, where tumor-driven bone loss supports disease growth; no safe intervention currently prevents its development or progression. Sigma anti-bonding calcium carbonate is an oyster shell–derived calcium carbonate in which weakened calcium–oxygen interactions lower the barrier to dissociation, making the mineral far more water-soluble and reactive than the conventional form. Here we characterized this modified calcium and tested whether it alters bone formation and myeloma progression in mice. The modified calcium was approximately 190-fold more soluble than conventional calcium carbonate and raised the pH of acetic acid far more rapidly. A single oral dose increased serum ionized calcium by more than 40% for 3 hours, and repeated dosing sustained this elevation, which reversed within 4 weeks of stopping treatment. The effect was greater in mice fed a vitamin D–insufficient diet, indicating absorption that does not require vitamin D. Treated mice showed enhanced osteoblast differentiation, greater trabecular bone mass, and higher serum bone formation markers, with elevated urinary calcium but no sign of kidney stones and reduced serum creatinine. In immunodeficient and immunocompetent mice transplanted with 5TGM1 myeloma cells, treatment begun 4 weeks before transplantation prevented engraftment or substantially delayed myeloma growth, increased bone formation, raised osteoblast numbers, lowered osteoclast numbers, and prolonged survival; treatment begun after transplantation had smaller but significant effects. These findings identify this bioavailable, bone-anabolic oral calcium as a candidate for preventing or delaying myeloma development and progression.

## Introduction

Multiple myeloma (MM) is the most common cancer of the bone marrow (BM)^1^ and remains a difficult-to-cure hematological malignancy due to resistance to chemo- and immuno-therapies. MM cells are terminally differentiated plasma cells that reside within the BM; these cells rely heavily on the surrounding microenvironment for survival and disease progression^2–5^. Previously, we and others hypothesized that improving bone formation may deter myeloma progression and tested novel and existing bone-anabolic drugs as a treatment for MM. However, they had only a marginal effect on MM progression^6–11^. Interestingly, the surviving mice from the treatments exhibited complete resistance to subsequent tumor rechallenge^12^. These results suggest that alterations in the BM microenvironment of the surviving mice may provide protective mechanisms against subsequent tumor engraftment.

Calcium is one of the most abundant minerals in the human body, 99% of which is stored in bones and teeth^13^. In the skeleton, calcium, along with phosphorus, provides structural integrity but also serves as a dynamic calcium-mineral reservoir, maintaining optimal blood calcium homeostasis^14,15^. High blood calcium concentrations stimulate calcitonin secretion from the thyroid gland, which induces osteoblast differentiation, matrix protein expression, and mineral nucleation. Conversely, low blood calcium concentrations stimulate the secretion of parathyroid hormone, which promotes bone resorption by osteoclasts and regulates systemic calcium balance^16,17^. This process is closely coordinated by endocrine regulators, including parathyroid hormone, vitamin D, and fibroblast growth factor 23, which integrate systemic calcium metabolism with local bone remodeling^18–20^. From a clinical perspective, the recommended daily calcium intake for adults is >1,000 mg, and adequate calcium intake is essential throughout life. In adults, calcium plays a pivotal role in bone remodeling and preservation, while insufficient intake or inadequate metabolism may lead to underserved mineralization, osteoporosis, and increased bone fracture susceptibility^21–23^. Thus, calcium serves as both a fundamental molecular mediator of bone formation and a key nutritional factor in maintaining skeletal integrity. Calcium and vitamin D supplementation is typically recommended as part of anti-aging or osteoporosis treatment, particularly for patients receiving osteoporosis medications^23^ and multiple myeloma therapy.

Sigma anti-bonding calcium carbonate (SAC) is extracted from oyster shells and then processed. Unlike conventional CaCO_3_, SAC exhibits reduced effective lattice stabilization due to weakened Ca–O interactions, thereby lowering the energetic barrier for dissociation and enhancing coordination with water. As a result, calcium in SAC exists in a more bioavailable, ion-like state under aqueous conditions^24^. Notably, SAC improved bone density by preventing bone restructuring mediated by osteocalcin, collagen, and minerals in ovariectomized rats^25^ and in patients with osteoporosis^26^. Due to its weak anti-bonding, SAC is presumed to be readily absorbed through the gut and produces ionized calcium in serum. In the bone environment, these elevated extracellular calcium ions may promote osteoblast differentiation, resulting in bone formation^27^. No side effects have been reported in either animal models^25^ or human studies^26^.

In the present study, we evaluated i) whether a SAC dietary supplement could elevate serum calcium ion levels, resulting in bone formation, and ii) whether pretreatment with SAC could delay the onset or progression of myeloma, while improving bone formation, in both immunodeficient and competent MM animal models.

## Results

### SAC ionizes calcium faster than CaCO_3_

SAC is a faster-reacting calcium ion and more soluble than normal calcium carbonate, CaCO ^24^. We tested SAC solubility in water (Fig. 1A) and its reactivity with acetic acid (CH_3_COOH) (Fig. 1B).

**Figure 1.**
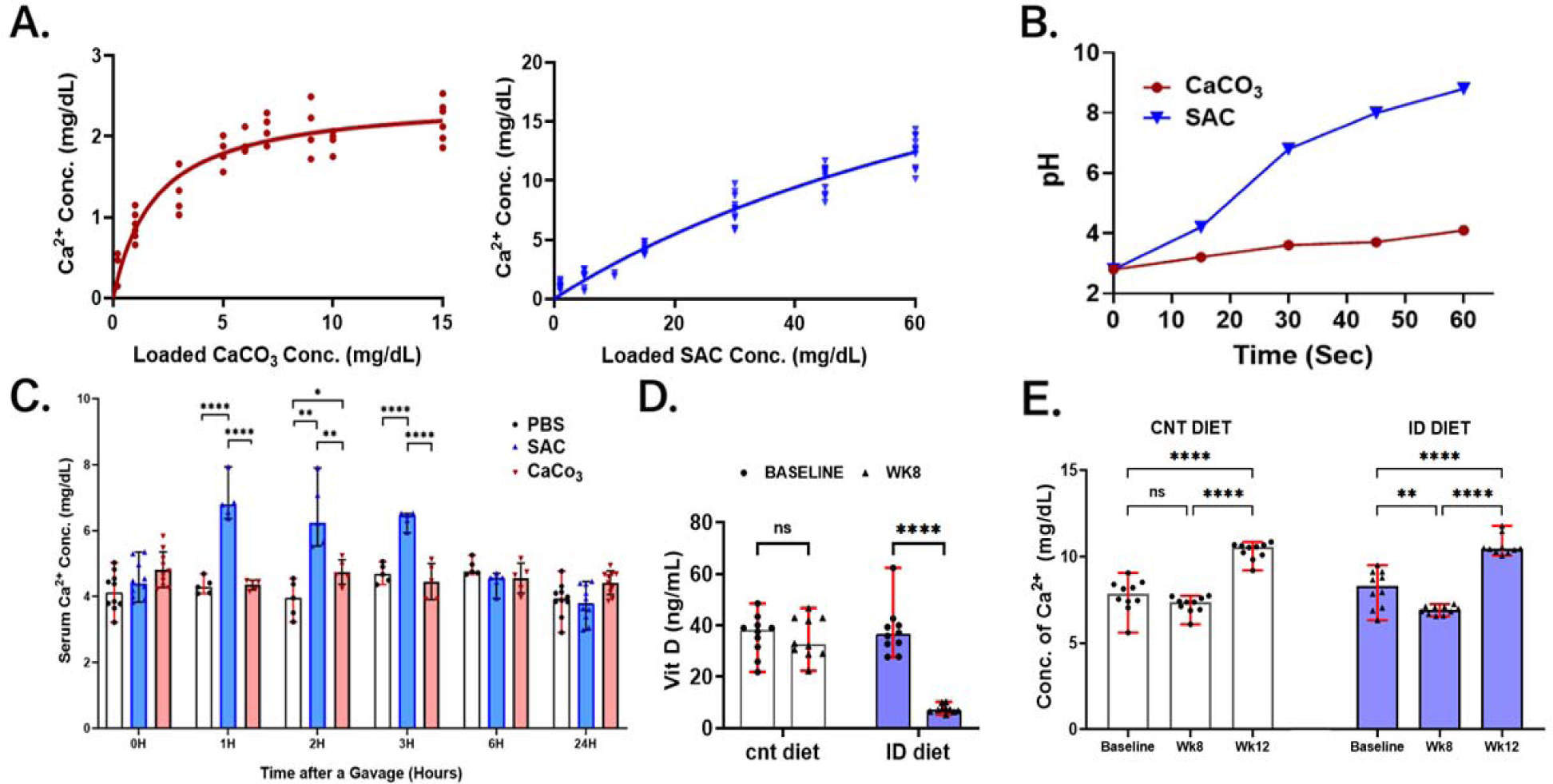
Characterization and *in vivo* pharmacokinetics of SAC: a novel calcium delivery model. (A) Solubility tests of SAC and CaCO_3_ in water. (B) pH changes of each SAC or CaCO_3_ mixture over 60 seconds. SAC (blue triangles) shows a rapid increase in pH over time compared to conventional calcium carbonate (red circles). (C) Serum calcium ion levels at indicated times after a single gavage of 150 µL of 1.5 mg/mL SAC, CaCO_3_, or 1X PBS. (D) Serum Vitamin D (ng/mL) levels measured under control (cnt diet) and Vitamin D-insufficient (ID diet) conditions. (E) Concentration of serum calcium (mg/dL in control diet and Vitamin D-deficient diet animals. Comparisons were analyzed with Student’s t-tests. * denotes p < 0.05; ** denotes p<0.01; **** denotes p < 0.0001; while non-significant was not shown.

For the solubility test, SAC or CaCO_3_ was diluted to concentrations of up to 15 mg/dL in distilled water. As shown in Fig. 1A, SAC shows a continuous increase in calcium ion concentration, while CaCO3 saturates at 5 mg/dL. We further added 30, 45, and 60 mg/dL of SAC to determine the saturation constant. At 60 mg/dL, SAC shows a slower elevation of calcium ions. Hyperbola least-squares fit analysis calculated that SAC showed a 13.88-fold increase in B_max_ (maximum binding: SAC, 34.29 vs. CaCO_3_, 2.47) and a 55.34-fold increase in K_d_ (dissociation constant: SAC, 105.7 vs. CaCO_3_, 1.91) compared to CaCO_3_. This indicates that SAC is 190 times more soluble than CaCO_3_.

To evaluate the reaction rate of SAC over CaCO_3_, we monitored pH changes for 60 seconds after adding either SAC or standard CaCO_3_ to acetic acid (Fig. 1B). The reaction of released calcium ions from SAC or standard CaCO_3_ with acetic acid increases the mixture’s pH over time. While the standard CaCO_3_ mixture increases the pH to 4.1, the SAC mixture rapidly raises the pH to 8.8 in 60 seconds. This rapid elevation in pH indicates that SAC reacts faster than CaCO_3_ (Fig. 1B).

### Oral SAC increases serum ionic calcium and bone formation

We then determined whether oral gavage of SAC, CaCO_3_, or PBS increased serum calcium ion levels in mice. For baseline, we collected sera a week before treatment. To ensure maximum uptake, we emptied the mouse stomach for 30 minutes before gavage by starvation. Thereafter, 228 µg in 150 µL of SAC, CaCO_3_, or 1X PBS was administered by oral gavage. To minimize the number of bleeds per mouse, mice were randomly assigned to 1-hour or 3-hour bleeding groups or to 2-hour or 6-hour bleeding groups. As shown in Fig. 1C, only SAC increased serum calcium ion concentration by >40%; this increase was maintained over 3 hours, then returned to baseline at 6 hours, while PBS- or CaCO_3_-treated groups showed no change.

To further characterize the long-term effects of SAC on the serum calcium ion level, we treated mice with PBS or SAC for two 4-week periods (4 weeks PBS or SAC, then split them into PBS or SAC groups in the subsequent 4-week period): i) PBS/PBS; ii) PBS/SAC; iii) SAC/PBS; and iv) SAC/SAC. Serum calcium ion levels increased during the first 4-week treatments with SAC (SAC/PBS and SAC/SAC); switching to PBS for the second 4-week treatment (SAC/PBS) restored normal calcium ion levels, while switching from PBS to SAC increased calcium ion levels (*p* = 0.013) (Fig. 2A).

**Figure 2.**
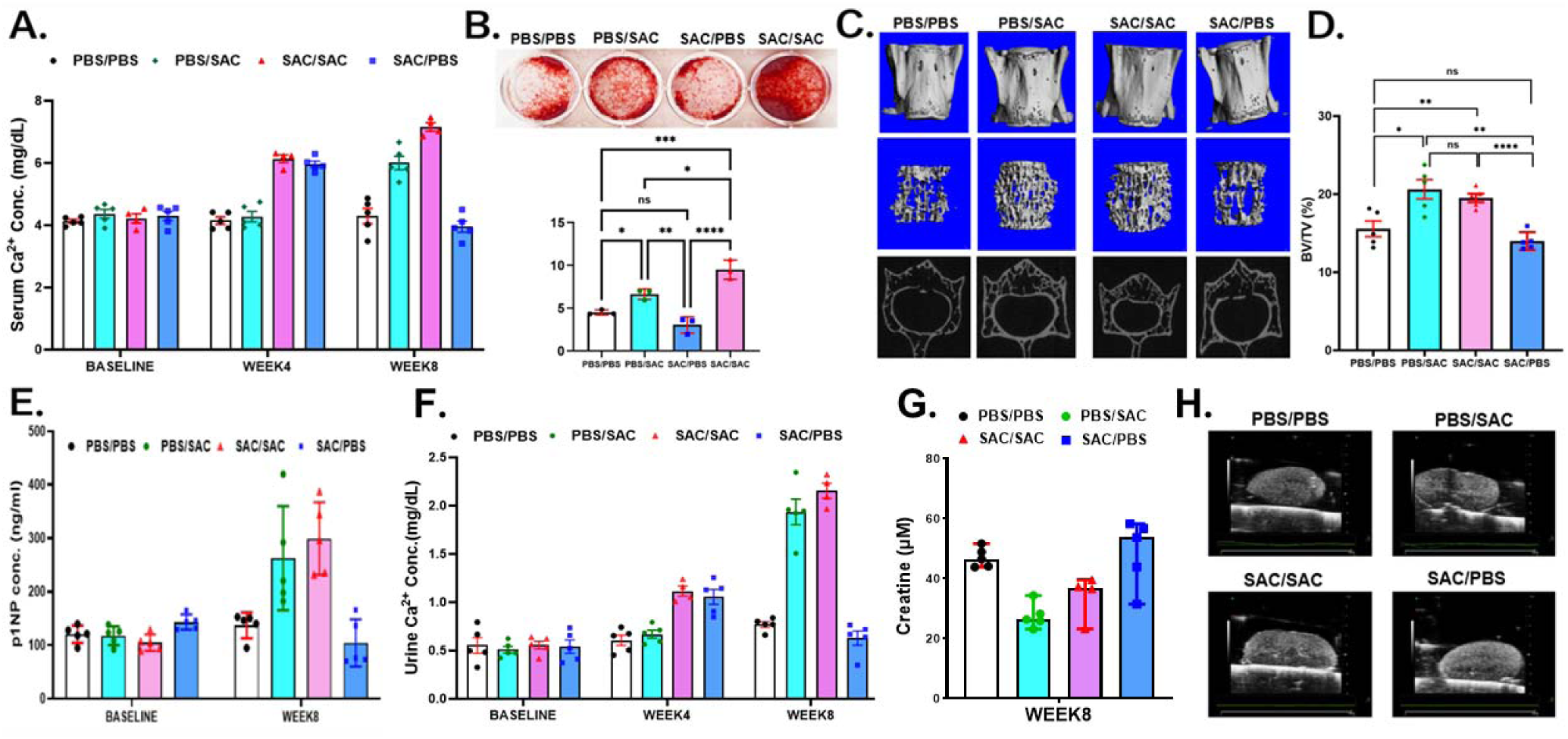
SAC promotes bone mineral density and bone turnover markers in mice. (A) Sigma anti-bonding calcium carbonate (SAC) elevates serum calcium levels based on treatments. The graph demonstrates serum calcium level changes at baseline, 4 weeks, and 8 weeks across 4 experimental conditions (PBS/PBS, PBS/SAC, SAC/SAC, SAC/PBS). (B) Alizarin Red S staining images showing calcium deposition after 8 weeks with different combination treatments, demonstrating different osteogenic potentials. (C) Cortical 3D, trabecular 3D, and 2D images. (D) Bone volume (BV/TV) of the SAC- and PBS-treated groups. (E) Serum bone anabolic marker p1NP ELISA. (F) Ionic calcium levels in the urine of SAC- and PBS-treated groups. (G) Creatinine concentration at week 8 across treatment groups. (H) *Ex vivo* kidney ultrasound images of the SAC- and PBS-treated groups. Comparisons were analyzed with Student’s t-tests. NS denotes not significant; * denotes p < 0.05; ** denotes p<0.01, **** denotes p < 0.0001

To test whether SAC absorption via the dietary route depends on Vitamin D, we induced a Vitamin D-insufficient state in mice by feeding them a Vitamin D-insufficient diet for 8 weeks. Serum Vitamin D levels were measured in the 8th week and demonstrated a highly significant reduction in metrics (*p* < 0.001), dropping by an average of 80.34% in the Vitamin D-insufficient diet group, while the control group remained statistically stable, experiencing a non-significant decrease of 3.30% (*p* = 0.70) (Fig. 1D). It also resulted in lower serum ionized calcium levels in the Vitamin D-insufficient diet group (*p* = 0.002, whilst *p* = 0.24 on control diet). We then administered SAC via gavage twice daily for 4 weeks. All SAC treatments in both the control and Vitamin D-insufficient diet groups increased serum ionized calcium levels. The increment (53.8%) in the Vitamin D-insufficient diet groups was higher than 43% in the control diet group (*p* = 0.0496, Fig. 1E).

To evaluate changes in osteogenic potential, we harvested bone marrow cells from the femurs and tibiae and performed *in vitro* osteoblast differentiation, followed by Alizarin Red S staining. As shown in Fig. 2B, the PBS/SAC and SAC/SAC groups showed increased mineralization, with a greater effect at 8 weeks in the SAC/SAC group. In contrast, in the SAC/PBS group, subsequent PBS treatment abolished the initial SAC-induced increase in osteogenic potential. Micro-CT analysis of the vertebra showed that SAC treatment for 4 weeks or longer (PBS/SAC and SAC/SAC) increased trabecular bone mass, as indicated by the bone volume/total volume (BV/TV) ratio. However, the 8-week SAC treatment (SAC/SAC) did not yield an additional increase in BV/TV over the 4-week treatment group. Consistent with the *in vitro* osteogenic potential results, the subsequent 4-week PBS treatment (SAC/PBS) reduced trabecular bone mass to a level comparable to that of the control group (PBS/PBS) (Figs. 2C and D). Likewise, the serum bone formation marker P1NP was also increased in the PBS/SAC and SAC/SAC groups (Fig. 2E). While high serum calcium levels increase bone formation, they may also cause kidney damage. We assessed urinary calcium levels in these mice and performed *ex vivo* ultrasound (FUJIFILM VisualSonics, Bothell, WA, USA) of their kidneys. SAC-treated mice had elevated urinary calcium (Fig. 2F) but no evidence of kidney stones (Fig. 2G), suggesting that the kidney excretes serum calcium without crystallization. We also measured serum creatinine in the 8-week-treated mice and found that SAC treatment decreases serum creatinine levels (Fig. 2H).

### SAC inhibits MM cell growth and induces bone formation in the 5TGM1-transplanted NSG mouse model

In MM bone disease (MMBD), MM cells interact with bone cells to promote bone resorption and MM cell growth. We hypothesize that SAC-mediated bone formation affects MM engraftment and/or growth in the MMBD mouse model. To test this hypothesis, we used an improved MMBD model that involves engrafting 5TGM1-*Luc* cells into NSG mice. This model allows for monitoring MM engraftment and development, which becomes evident at 14 days post-transplant, using longitudinal, weekly *in vivo* bioluminescence imaging^28^.

To maximize the bone-anabolic effect of SAC, we designed the study with 3 groups: i) 4 weeks pretreatment before transplant (TbT); ii) treatment 2 weeks after transplant (TaT); and iii) PBS treatment for the study period, as described in Fig. 3A. While PBS-treated mice exhibited MM engraftment and disease progression and 100% mortality at 45 days post-transplant, both SAC treatments (TbT and TaT) significantly delayed tumor growth (Fig. 3B and C). Furthermore, the bioluminescence signals from TbT mice decreased over time (Fig. 3C). We performed the Mantel–Cox test to determine whether SAC pretreatment (TbT) or posttreatment (TaT) affected survival and found that both treatments significantly extended the animals’ median survival (undefined, *p* = 0.0007 for the TbT group and 45 days, *p* = 0.0437 for the TaT group) compared to PBS (39 days) (Fig. 3D).

**Figure 3.**
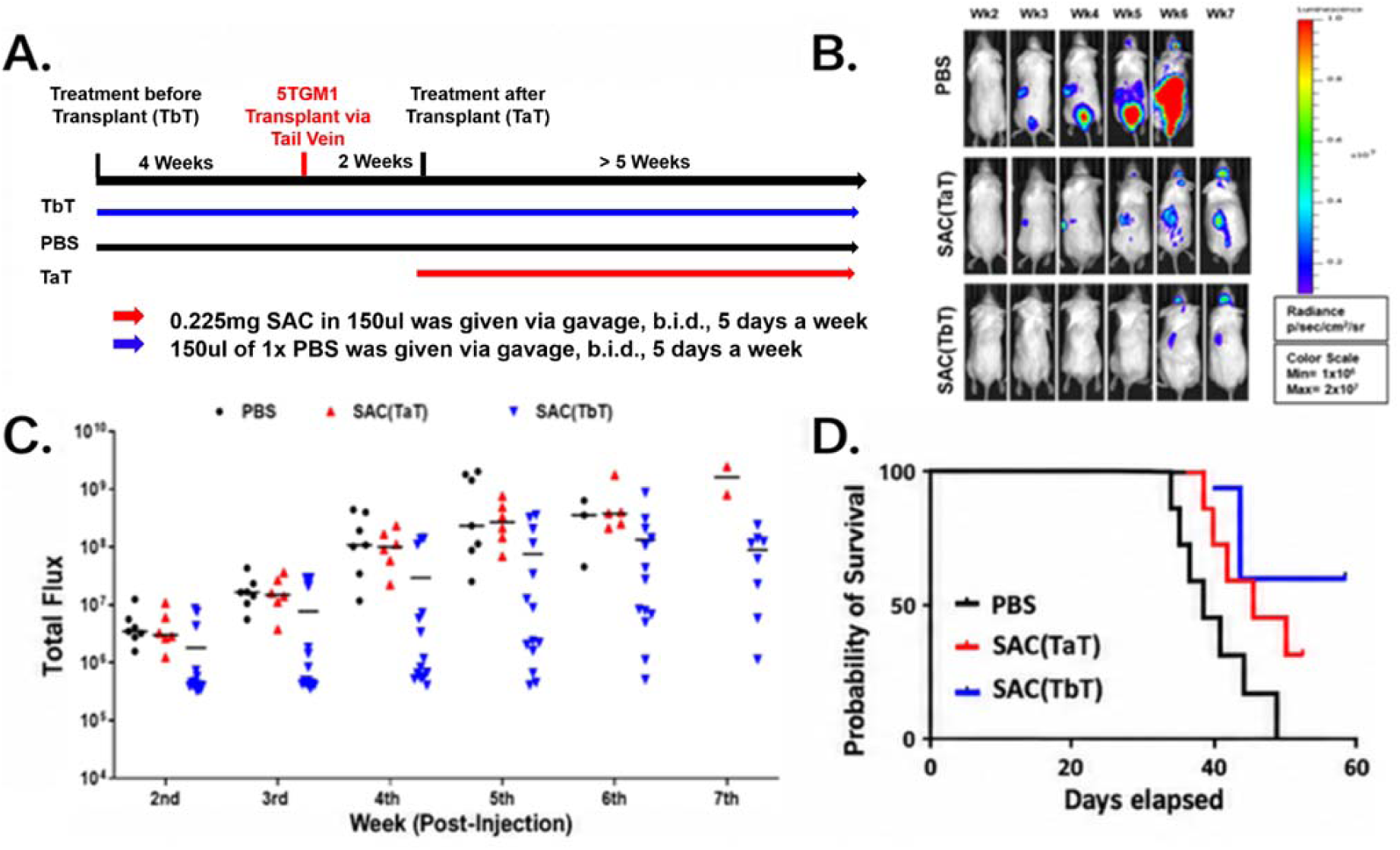
SAC inhibits MM cells and extends mouse survival. (A) Experimental design. (B) Representative bioluminescence images of PBS-treated vs SAC-treated 5TGM-1-luc transplanted NSG mice. All bioluminescent images were normalized to a single scale, as shown in the box. (C) From the weekly bioluminescent image, a total flux value was obtained for the whole body of the mouse. All total flux values (black dot, or red and blue triangles; PBS- or SAC-treated) with median (black line) were shown here. (D) Both TaT and TbT mice experience delayed or no MM progression. Mouse survival was scored daily. When the mouse met the endpoint criteria, it was euthanized. Survival curves were plotted from 7 mice from PBS (black line), 7 mice from SAC (TaT) (red line), and 14 mice from SAC groups (TbT) (blue line). Median survival times were 39 days for PBS, 45 days for SAC-TaT, and undefined for SAC-TbT groups. The Mantel–Cox test showed p = 0.0437 for PBS vs SAC (TaT) and p = 0.0007 for PBS vs SAC (TbT). SAC: sigma anti-bonding calcium carbonate.

To evaluate the skeletons of these mice, we measured spine bone mineral density in SAC-treated (TbT and TaT) and control mice using a PIXImus densitometer. Bone mineral density at the spine increased significantly in TaT (*p* = 0.003) and TbT (*p* = 0.030) mice compared with controls (Fig. 4A). We also performed bone histomorphometry analyses using micro-CT. SAC-treated groups had significantly higher bone volume density (TaT, *p* < 0.003; TbT, *p* < 0.018) than the control group (Fig. 4B); trabecular thickness also increased significantly (TaT, *p* < 0.008; TbT, *p* < 0.022) compared to the control group (Fig. 4C). To assess the number of active osteoclasts and osteoblasts, the third and fourth lumbar vertebrae from PBS and TbT mice were processed and prepared for plastic-embedded slides, then stained with tartrate-resistant acid phosphatase to count osteoclasts and osteoblasts. We found significantly fewer osteoclasts (*p* < 0.0001) and significantly more osteoblasts (*p* < 0.003) in TbT mice than in controls (Fig. 4D and E).

**Figure 4.**
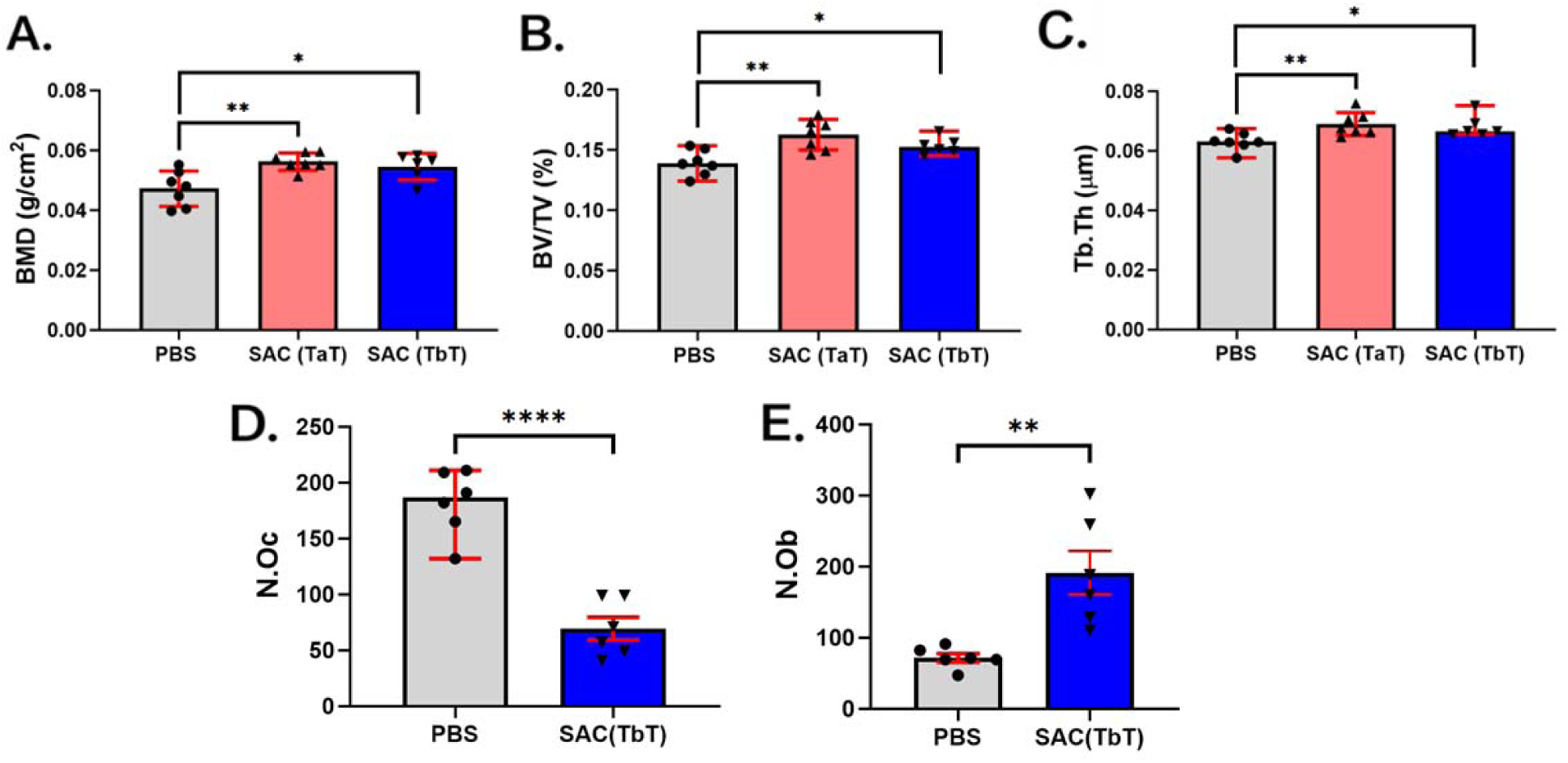
Bone formation increases in both SAC-treated 5TGM1-*Luc*-transplanted NSG mice. Treated and non-treated 5TGM1-*Luc*-transplanted NSG mouse vertebrae were scanned for DEXA using PIXImus. **(A)** Bone mineral density (BMD) changes were shown in the whole spine of SAC-TaT (mice surviving >6 wks) and SAC-TbT (mice surviving >7 wks) mice versus PBS. The mouse vertebrae were extracted/scanned by microCT. Myeloma-associated bone lesions were assessed at the trabecular regions of total lumbar vertebrae (L1–L6). **(B)** bone volume to total volume (BV/TV) and **(C)** trabecular thickness (Tb.Th) are shown. The L3 and L4 of PBS and SAC (TbT) were further prepared for tissue section and stained with TRAP. Osteoblasts and osteoclasts were counted. **(D)** Osteoclast numbers (N.Oc) in PBS-treated vs SAC-TbT are shown. **(E)** Osteoblast numbers (N.Ob) in PBS-treated vs SAC-TbT are shown. Comparisons were analyzed with Student’s t-tests. * denotes p < 0.05, ** denotes p < 0.01, **** denotes p < 0.0001.

### SAC pretreatment delays MM progression and induces bone formation in the 5TGM1-transplanted C57BL/KaLwRij mouse model

The 5T MM mouse models have been instrumental in defining MM potential, studying MMBD biology, and examining novel therapeutics^29–32^. Among them, 5TGM1-transplanted cells exhibit short dormancy in the C57BL/KaLwRij model, and animals develop a tumor burden with pronounced osteolytic lesions^6,29^. While this is a well-recognized MMBD model with immune competence, animals show significant variation in bone lesions and MM development compared with the NSG MM model^28^.

To evaluate the effects of SAC on MM growth in the well-established immune-competent MMBD model, C57BL/KaLwRij mice were treated with SAC or PBS starting 4 weeks before MM cell transplant (TbT or PBS). Then, 1 × 10^6^ 5TGM1-*Luc* cells per mouse were injected into both groups. Survival was scored daily for 9 weeks. Survival increased significantly in the TbT mice compared to controls (*p* = 0.0007); the median survival time for TbT mice was undefined but was 46 days in the control group (Fig. 5A). Bone mineral density increased in the TbT mice compared to the control group (*p* = 0.0429) (Fig. 5B), and bone histomorphometry analyses showed increases in the BV/TV and trabecular thickness in the spines of TbT mice (*p* = 0.001 and *p* = 0.0001, respectively) (Fig. 5C and D). We also observed that SAC significantly decreased the number of osteoclasts and significantly increased the number of osteoblasts (*p* = 0.0001 for both) (Fig. 5E). These results provide strong evidence that SAC TbT treatment delays MM progression and improves bone formation, even in the conventional MMBD mouse model.

**Figure 5.**
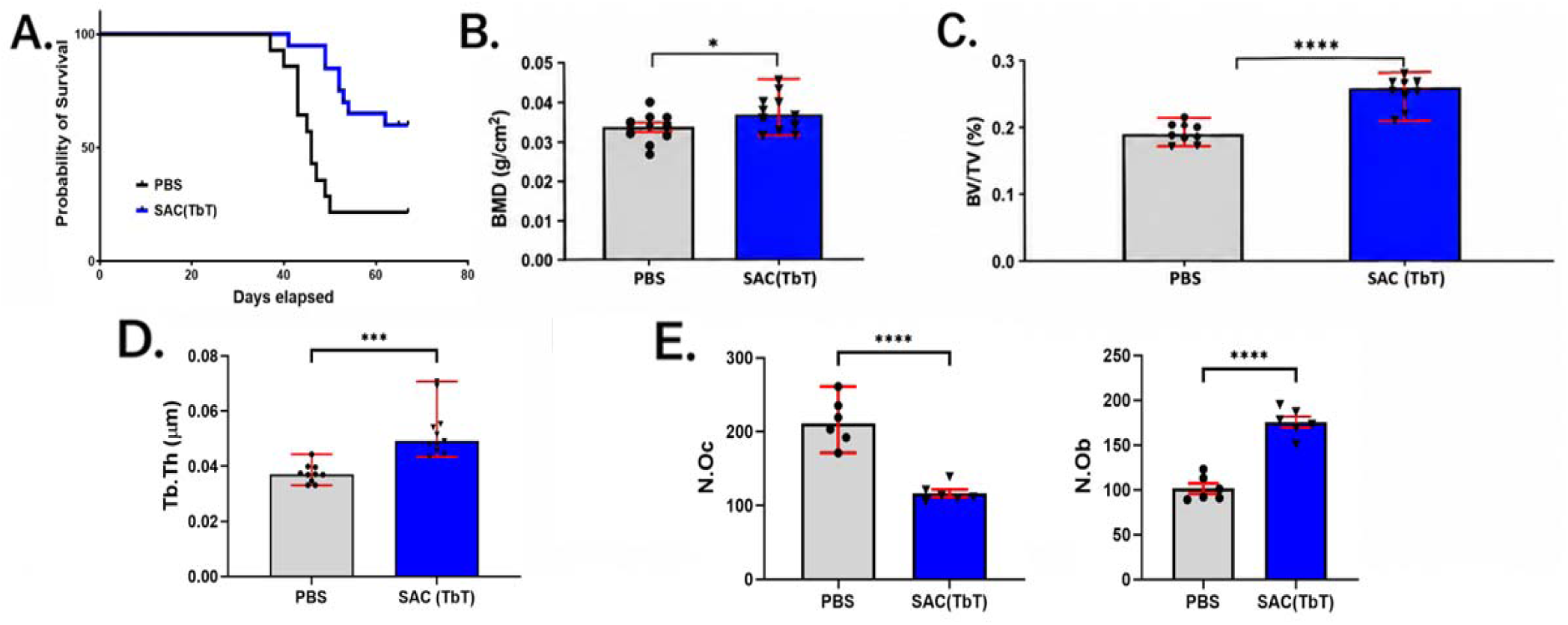
SAC-TbT delays MM progression and bone formation in 5TGM1-*Luc*-transplanted C57/KaLwRij mice. (A) Survival curves for 14 mice from PBS (black line) and 20 mice from SAC groups (TbT) (blue line) (p=0.0017). Median survival times were 39 days for PBS and undefined for SAC-TbT-treated mice. Vertebrae were extracted from the frozen body and analyzed with DEXA and microCT. (B) Bone mineral density (BMD), (C) BV/TV, and (D) Tb.Th in spines of SAC- and PBS-treated mice. Number of osteoblasts increased in SAC-treated C57/KaLwRij mice, while osteoclasts decreased. (E) Osteoclast numbers (N.Oc) and osteoblast numbers (N.Ob) in PBS-treated vs SAC-treated (TbT) mouse spines. Comparisons were analyzed with Student’s t-tests. * denotes p < 0.05. *** denotes p < 0.001. **** denotes p < 0.0001. MM: multiple myeloma. SAC: sigma anti-bonding calcium carbonate.

## Discussion

Calcium is the principal cation of bone. While intracellular calcium plays a signaling role in many cell types, including bone cells^27^, bone is the major source/contributor to extracellular (serum) calcium homeostasis. Calcium enters the body primarily through dietary intake. It is mainly absorbed in the duodenum and portions of the small intestine through an active vitamin D-dependent mechanism, with a smaller portion absorbed through passive diffusion. The active processes are saturable. With high calcium intake, passive uptake via diffusion predominates^33^. Calcium is filtered by the kidneys, and approximately 99% of the filtered calcium (about 60% of total plasma calcium not bound to proteins) is reabsorbed in the renal tubules via hormone-dependent active transport. The exchange between extracellular calcium and bone mineral may occur via rapid non-hormonal mechanisms or slower hormone-mediated processes^33^. Cumulative evidence has shown that increasing calcium intake reduces fracture rates and prevents bone loss^34,35^. At the same time, high calcium intake may lead to hypercalcemia or kidney stone concerns. No clinical issue associated with high calcium intake has been reported^35^.

SAC forms Ca(H_2_O)_6_^2+^ in aqueous solution and becomes more soluble^24^. Our results showed that SAC has exceptional water solubility and greater potency compared to an equimolar concentration of conventional calcium carbonate. We observed that a single dose of SAC administered by oral gavage transiently elevated serum calcium ion levels over 2 hours (levels returned to baseline after 6 hours), whereas conventional calcium carbonate or PBS did not have the same effect. Our results in diet-induced Vitamin D-insufficient mice suggest that the body absorbs SAC via passive uptake (independent of Vitamin D). Continuous SAC intake sustained elevated serum calcium levels, while discontinuation of SAC restored serum calcium levels to the normal range within 4 weeks. These results suggest that SAC may be a potential therapeutic intervention for patients with Vitamin D insufficiency.

Increased extracellular calcium ions in the bone microenvironment can promote osteoblast differentiation and stimulate bone formation^27^. Earlier and concurrent studies by other groups found that SAC increased bone density by preventing osteocalcin-mediated bone resorption in ovariectomized mice^36^ and rats^25^. Our study showed that 4-week SAC treatment was sufficient to maintain elevated serum calcium ion, but this effect was abolished 4 weeks after discontinuation. In addition, we found a significant increase in bone formation. Although hypercalcemia can affect kidney function, we observed elevated urinary ionic calcium with reduced serum creatinine and no sign of kidney stones.

MM is a hematologic malignancy of terminally differentiated plasma cells that reside within the BM and rely heavily on the surrounding microenvironment for their survival and disease progression^2–5^. As we and others tested novel or existing bone-anabolic drugs as treatments for MM, these studies showed only marginal effects on myeloma progression^6–11^. Given the considerable effects of SAC on bone formation and its safety profile, we tested a novel regimen in which MM cells (5TGM1-*Luc*) were engrafted via the tail vein after 4 weeks of pretreatment with SAC (TbT). While the conventional treatment regimen (TaT) exhibited marginal effects on myeloma progression (20% survival), animals in the TbT group showed 60% survival at 7 weeks post-transplantation and exhibited an exceptional delay in MM progression. All SAC-treated (both TbT and TaT) mice exhibited significant bone formation, with a significant increase in the number of osteoblasts and a decrease in the number of osteoclasts. We found similar results in the conventional MMBD model.

In conclusion, we summarize our major findings alongside previous discoveries to describe how SAC works against MM in Fig. 6. Briefly, SAC markedly increased serum calcium ion levels via a Vitamin D-independent pathway, leading to bone formation without causing renal impairment, and its effects persist only during SAC uptake. Furthermore, SAC showed remarkable anti-MM effects, with significant bone formation when administered before the 5TGM1-*Luc* transplantation. These results suggest that SAC could serve as a novel preventive or adjunctive treatment for MM development or recurrence, or for associated bone complications.

**Figure 6.**
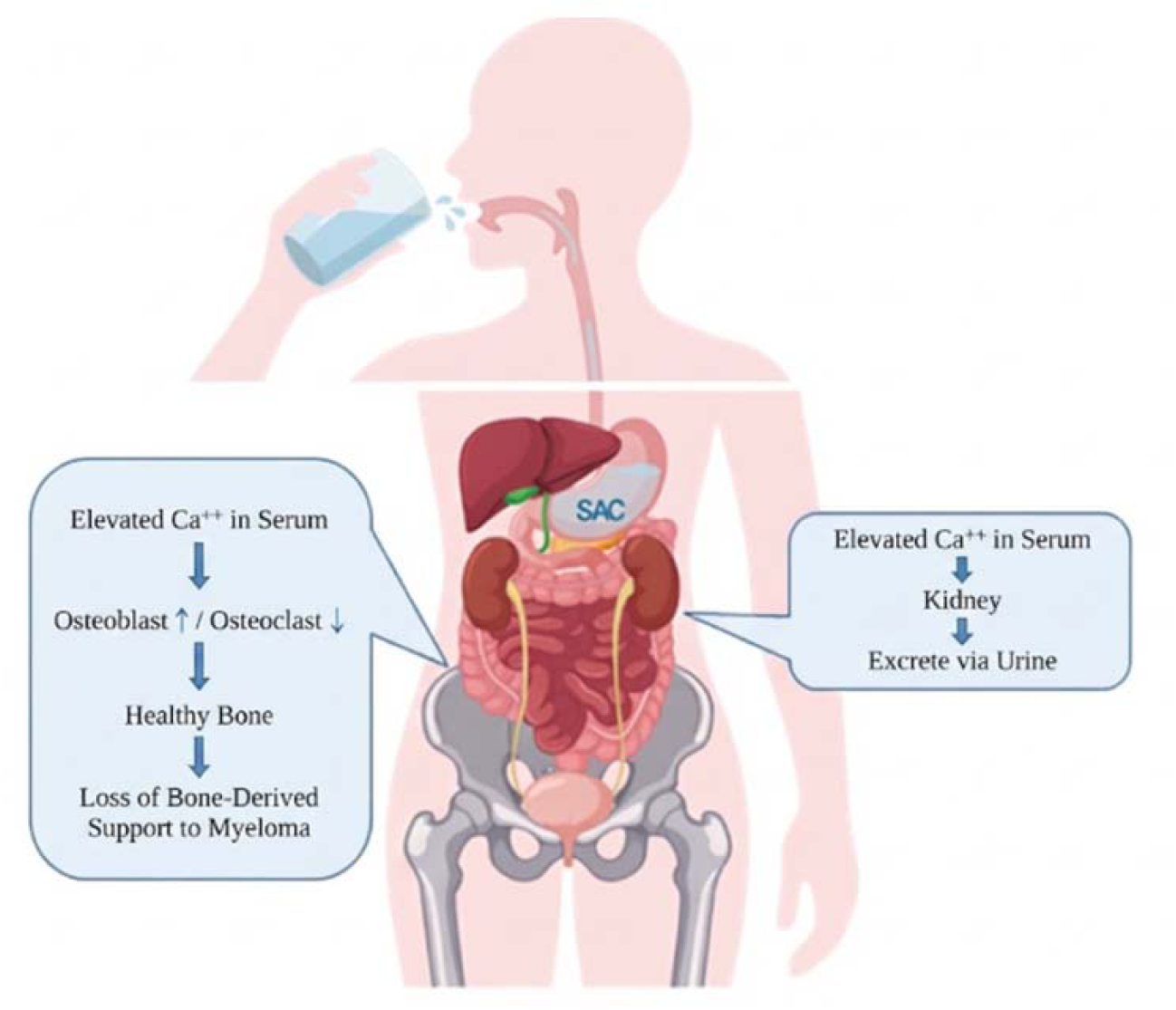
Summary of Our Findings.

## Materials and Methods

### Study design

This study had three objectives: (i) to compare the aqueous solubility and acid reactivity of SAC with those of conventional CaCO_3_; (ii) to determine whether oral SAC raises serum ionized calcium and promotes bone formation in normal and vitamin D–insufficient mice, and whether it affects renal function; and (iii) to determine whether SAC given before (TbT) or after (TaT) transplantation of 5TGM1-*Luc* myeloma cells delays myeloma engraftment or progression and improves bone parameters in immunodeficient (NSG) and immunocompetent (C57BL/KaLwRij) mice.

### Test materials

SAC solution (1.5 mg/mL) was obtained from the Calcium & Bone Health Institute (Coquitlam, British Columbia, Canada). It contains 1.5 mg SAC Calcium, 8 mg Selenium, 0.05 mg Magnesium, 0.02 mg Zinc in 1 ml SAC solution. We used CaCO_3_ powder (Cat. C1074, Spectrum Chemical MFG Corp., Gardena, CA, USA)

### Solubility test

SAC and CaCO_3_ were prepared and diluted to concentrations of 1, 3, 5, 6, 10, and 15 mg/dL in distilled water. All solutions were mixed/incubated for >40 minutes, then centrifuged at 10,000 × *g* for 5 minutes to remove any undissolved material. Only 5 µl of the supernatant was subjected to a colorimetric calcium assay (Metallo Assay, Funakoshi, Cat. No. CA31ME, Tokyo, Japan). Additionally, SAC at 30, 45, and 60 mg/dL was added to the assay to determine the saturation constant. The B_max_ and K_d_ of SAC and CaCO_3_ were calculated by least-squares fit to a hyperbolic equation using Prism 11 (GraphPad Software Inc., San Diego, CA, USA).

### Reaction rate test

The reaction rates of SAC and calcium carbonate were measured using a digital pH meter (Mettler Toledo, Five Easy Plus, FP20, Columbus, Ohio, USA) based on the following chemical equation:

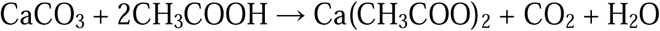

Equal masses of SAC or calcium carbonate were added to the beaker with 500 mL of 0.1 M acetic acid solution (CH_3_COOH) to reach a theoretical maximum calcium ion concentration of 0.2250 mol/L. To ensure a uniform reaction, the solutions were stirred at a constant speed using a magnetic stirrer. The pH was assessed at 15-second intervals from 0 to 60 seconds.

### Animals and ethics statement

All mouse strains were housed and bred at the University of Arkansas for Medical Sciences (UAMS) Animal Facility. All animal procedures were approved by the UAMS Institutional Animal Care and Use Committee (Protocol no. IPROTO202400000149) and were conducted as per the National Institutes of Health (NIH) Guide for the Care and Use of Laboratory Animals. Mice were kept on a 12 h light/12 h dark cycle. Room temperature was maintained at 24–26 °C. Food pellets and water were sterilized and provided as needed. All animal studies used equal numbers of males and females.

### SAC, CaCO_3_, and PBS treatments

NOD.Cg-*Prkdc^scid^ Il2rg^tm1Wjl^*/SzJ (NSG) or C57BL/KaLwRij mice (males and females, 8 to 12 weeks old) received 150 µl of 1.5 mg/mL SAC or CaCO_3_ or 1X PBS by oral gavage twice daily with 6-hour gap, 5 consecutive days/week for 4 weeks before 5TGM1-*Luc* cell transplantation (TbT) or 2 weeks after transplantation (TaT), while PBS served as the control.

### Vitamin D–insufficient mouse model

Seven- or eight-week-old C57/BL6 mice were switched to the Vitamin D-insufficient diet (D08090903, Research Diets Inc.) or a control diet (D10001, AIN-76A) (n=10 per dietary group). During the experimental period, mice had ad libitum access to water and food in a room with controlled temperature. Mice were housed in standard cages and maintained on a standard 12-h-light/-dark cycle. Mice were maintained on the custom diets till the end of the experiment.

### Quantification of serum vitamin D concentration

At different time points during the trial, blood was collected by retro-orbital bleeding from both test and control diet groups. Following collection, blood was separated into serum and used for biochemical assays. To measure vitamin D levels, we used a mouse 25-hydroxyvitamin D3 (25 HVD3) ELISA kit (DC-CSB-E08099m_96T, CUSABIO, Houston, TX, USA, sensitivity 23.5 pg/ml) according to the manufacturer’s instructions.

### Quantification of calcium concentration

By retro-orbital bleeding, >20 µl of blood was collected and separated by centrifugation at 6,000 × *g* for 5 min. The plasma was transferred into other vials and stored at −80 °C for the assay. Serum calcium ion levels were analyzed using a colorimetric calcium assay kit (Metallo Assay, Funakoshi, Cat. No. CA31ME, Tokyo, Japan) according to the manufacturer’s instructions.

### P1NP measurement

We collected 100 µl of blood by retro-orbital bleeding and separated it by centrifugation at 9,300 × *g* for 5 min. The plasma was transferred into different vials and stored at −80 °C for the assay. Serum P1NP levels were determined with a mouse P1NP ELISA kit (R&D Systems, Minneapolis, MN, USA) according to the manufacturer’s instructions.

### Osteoblast differentiation

Bone marrow cells were obtained using a 28-gauge syringe by flushing the long bones of treated mice with α-MEM medium. Cells were collected from 5 mice in each group (PBS/PBS, PBS/SAC, SAC/PBS, and SAC/SAC), pooled, and cultured at 37 °C in an atmosphere of 5% CO_2_/95% air with 20% fetal bovine serum, 1% penicillin-streptomycin, and 50 μg/mL ascorbic acid (Sigma-Aldrich) in 10-cm culture dishes for 5 days. The medium was replaced every 3 days. Adherent (stromal) cells were re-plated in 12-well tissue culture plates at 2 × 10^5^ cells per well with 10% fetal bovine serum, 1% penicillin-streptomycin, 50 μg/mL ascorbic acid, and 10 mM β-glycerophosphate (Sigma-Aldrich) and incubated at 37 °C in an atmosphere of 5% CO_2_/95% air for 16 days to assess mineralization. The mineralized matrix was stained with 40 mM Alizarin red solution, according to the manufacturer’s instructions (Sigma-Aldrich). Alizarin red staining was quantified by extracting the dye with 10% acetylpyridinium chloride or 10% acetic acid and measuring the absorbance at 562 nm. For all assays, cells were plated in triplicate.

### Cell line and culture

5TGM1-*Luc* cells were obtained from the University of Texas Health Science Center, San Antonio, and maintained in RPMI 1640 medium containing 15% fetal bovine serum, 1X penicillin/streptomycin, and 1X GlutaMAX (Gibco, Life Technologies, U.K.) at 37 °C in an atmosphere of 5% CO_2_/95% air. 5TGM1-*Luc* cells were maintained in a range of 0.5–2 × 10^6^ cells/mL. Before being transplanted into mice, cells were washed 3 times with PBS and counted using a Cellometer Mini (Nexcelom, Lawrence, MA) with the trypan blue exclusion method.

### Multiple myeloma allograft mouse models

NSG and C57BL/KaLwRij mice were housed and bred at the UAMS animal facility. NSG and C57BL/KaLwRij mice (male and female, 8 to 12 weeks old) were inoculated intravenously with 10^6^ 5TGM1-*Luc* cells suspended in 100 µl PBS or with PBS alone. Myeloma cell engraftment was confirmed 2 weeks later with bioluminescence imaging.

### In vivo bioluminescence imaging and survival study

Tumor burden was monitored weekly from weeks 1 to 8 after 5TGM1-*Luc* transplantation with an *in vivo* Bioluminescence IVIS Imaging System 200 Series (PerkinElmer, Waltham, MA, USA). Mice were anesthetized using isoflurane, and images were acquired 10 minutes after intraperitoneal injection of 1.5 mg XenoLight D-luciferin potassium salt (PerkinElmer). All images were acquired using auto exposure. Using Living Image 4.7.4 software, a region of interest was generated to cover the whole body, and the total flux (p/s) was obtained. Mouse survival was scored daily. When a mouse met the endpoint criteria (i.e., hind limb paralysis, >20% body weight loss, body condition scoring <2), it was euthanized. The study was terminated 1 week after all mice in the PBS group died due to MM progression.

### Lumbar vertebral bone mineral density and bone histomorphometry

Mice were euthanized when the endpoint criteria were met (i.e., hindlimb paralysis, >20% body weight loss, and body condition scoring <2) or the study ended (8 weeks post-transplant). We stored euthanized mice in the freezer for at least 5 days. To compare mice with similar MMBD levels, we selected mice that survived ≥6 weeks. Mice were thawed at room temperature, and the spines were extracted. The bone mineral density of the vertebrae was measured using a PIXImus bone densitometer with on-board PIXImus software (GE Lunar, Madison, WI, USA), adjusted for body weight. Whole lumbar vertebrae (L1–L6) were scanned with a microCT40 (SCANCO Medical, Bassersdorf, Switzerland). For histomorphometric analyses, trabecular bone regions were selected to assess total volume and structural appearance in the lumbar vertebrae. The analyses provided information on the main histomorphometric parameters, including bone volume (BV/TV, %), trabecular thickness (Tb.Th), number (Tb.N), and spacing (Tb.Sp).

### Bone histological and cytochemical analyses

The extracted spines were fixed with Millonig’s fixative. Samples were sectioned at 5 or 7 μm and stained with hematoxylin and eosin. Serial spine sections were stained with hematoxylin and eosin and for tartrate-resistant acid phosphatase activity, a marker of osteoclasts in mouse bone *in vivo*.

### Statistical analysis

Statistical analyses were conducted with the Mantel–Cox test and Student’s *t*-test. A *p*-value of <0.05 was considered significant. GraphPad Prism 11 (GraphPad Software Inc., San Diego, CA, USA) and SigmaPlot 13.0 (Systat Software Inc, San Jose, CA, USA) were used for all statistical analyses.

## Data availability

The data presented in this study are available from the corresponding author upon request.

## Acknowledgements

This work was supported by grants from the Calcium & Bone Health Institute (CBHI CRA C23992, to D.Y.), the UAMS Winthrop P. Rockefeller Cancer Institute Seed of Sciences (to D.Y.), and the National Institute of General Medical Sciences of the NIH (P20GM125503, to D.Y.). This work was also supported in part by Merit Review Award I01 BX006235 (to F.Z.) from the U.S. Department of Veterans Affairs Biomedical Laboratory Research and Development Service; the contents do not represent the views of the U.S. Department of Veterans Affairs or the United States Government. Additional support came from the National Cancer Institute (2R01CA236814-06A1 and U54CA272691-01, to F.Z.), the Paula and Rodger Riney Foundation (to F.Z.), the Myeloma Solution Fund (to F.Z., Q.Y. and R.O.), Blood Cancer United (6702-25, to F.Z.), the International Myeloma Society (2TRAG2\100008, to F.Z.) and the UAMS Winthrop P. Rockefeller Cancer Institute Fund (to F.Z.).

CBHI, a non-profit organization, funded this study and supplied SAC solutions but did not make any decision regarding investigation or publication.

The authors thank the Science Communication Group at the University of Arkansas for Medical Sciences for editorial assistance, the Bone Histology and Imaging Core for imaging and analyses, and Ben Liu and Danna Kim for creating the graphics in Fig. 6.

